# Reanalysis of data from ten years of koala chlamydia vaccine trials reveals inherent bias and a lack of protective efficacy

**DOI:** 10.64898/2026.09.02.748764

**Authors:** Keith J. Chappell, Michaela D. J. Blyton

## Abstract

Australia’s iconic koalas are facing multiple threats from habitat loss, climate change and pervasive chlamydial disease. A safe and effective vaccine to protect koalas against chlamydia would be a welcome veterinary intervention to prevent this devastating disease which causes blindness, renal and reproductive pathology, infertility and mortality. However, over a decade of vaccine development and research trials has produced conflicting results, which vary from no observable indication of protective efficacy to a 64% reduction in chlamydia associated mortality. Given this discrepancy we conducted an independent analysis of data underpinning the highly cited publication demonstrating vaccine efficacy that was the basis for a permit for limited use in koalas. In doing so, we identified a substantial error in the way the data was encoded for analysis, which resulted in survival from birth to time of vaccination incorrectly being attributed to vaccination. After correction for this error, we show using the exact methods of the original authors that there was no significant difference in the survival or disease rate between vaccinated and unvaccinated koalas. Furthermore, we identified a substantial source of bias whereby animals with disease at their first health assessment were not excluded from the study and as those animals were generally not vaccinated, they were almost six times more likely to be included in the control group. Reanalysis including only koalas that were healthy on initial assessment and for which follow-up health assessments were performed, showed that vaccinated individuals were equally likely to contract disease or to die with disease; 18 of 143 vaccinated koalas (12.6%) developed disease and 3 (2.1%) died, compared to 63 of 439 control animals (14.4%) that developed disease and 11 (2.5%) that died. This study highlights the importance of randomised, observer blinded, placebo-controlled studies in veterinary settings to uphold scientific rigor and prevent stratification bias.

**Visual abstract:** 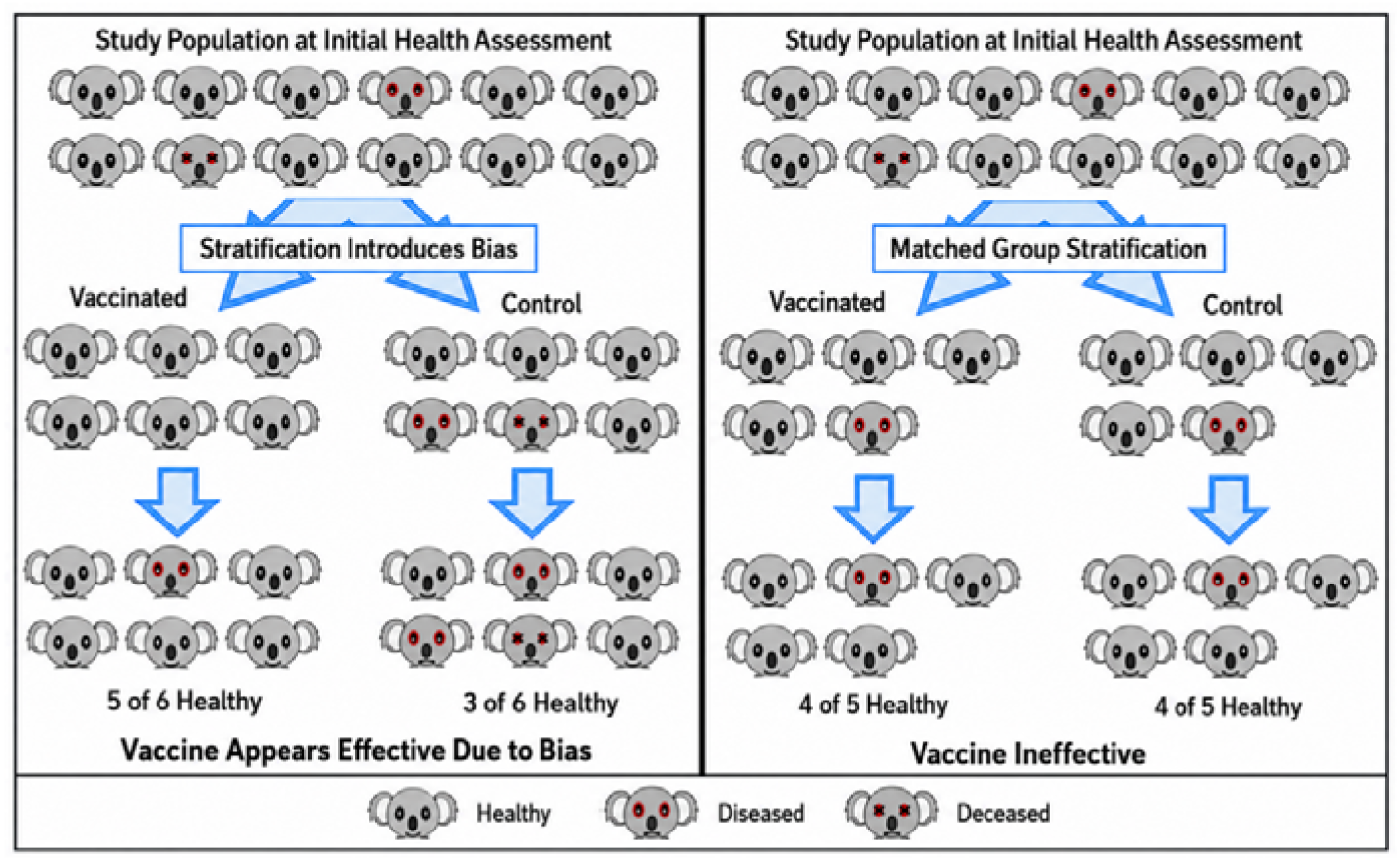

## Introduction

Koalas (*Phascolarctos cinereus*) were officially listed as endangered in the states of Queensland and New South Wales by the Australian Government in 2021 due to severe population declines throughout much of their northern geographic range [1]. This sparked concerted efforts by the Australian federal and state governments to develop, fund and implement management programs to conserve this iconic species. Along with habitat loss and climate change, infertility and mortality caused by *Chlamydia pecorum* infection has been identified as a major factor determining koala population viability [1]. While climate change and habitat loss are complex and political issues which require societal change, combatting disease in koalas is a more tractable area for rapid positive impact through veterinary interventions and active population management.

In September 2025, a koala chlamydia vaccine, Klavax^TM^ (Tréidlia Ltd) was granted a minor use permit in koalas by the Australian Pesticides and Veterinary Medicines Authority (APVMA). In what has been heralded as a breakthrough for koala conservation, the vaccine was widely promoted as safe and able to protect against chlamydia symptoms and disease associated mortality [2, 3]. However, conflicting results have emerged regarding the vaccine’s efficacy that calls into question whether it will be able to provide the much-anticipated means of combating chlamydia.

Phillips et al., [2] reported that vaccination provided significant protection against chlamydial disease and a 64% reduction in chlamydia associated mortality compared non vaccinated koalas from the same population. In contrast, Simpson et al. [4], reported that vaccination provided no protection against chlamydial disease. The trial performed by Simpson et al. [4], utilised a randomised, observer-blinded, placebo-controlled design, in which 102 wild koalas received the vaccine and 94 received a placebo. Following dosing, koalas were released into their natural environment and recaptured at 2, 6 and 12 months post-vaccination for clinical assessment by a veterinarian who was not aware of the koalas’ vaccination status. Conversely, the publication by Phillips et al. [2], is an aggregate analysis of five separate vaccine trials conducted over a ten year period [5-9] along with long term monitoring data on the populations in which the studies were performed.

Randomised, observer-blinded, placebo-controlled vaccine trials, as carried out by Simpson et al. [4], are the industry standard to determine the level of protection afforded against infection under natural conditions and are recommended by APVMA guidelines for licensure of new products [10]. This is because such trials remove the potential for the introduction of bias into the study design and ensure that any observed benefit of the vaccine cannot be the result of other factors such as pre-vaccination health, age of vaccination or treatment/intervention protocol. In other words, it ensures that the only difference between the treatment and control groups is that the treatment group received the vaccine. While such trials can be logistically challenging in wildlife species, Simpson et al. [4] demonstrates that they are possible and very valuable given their substantial advantages.

Nonetheless, the five vaccine trials combined in the study by Phillips et al. [2], cover an extended period of 10 years and included a large sample size of 680 koalas (although only 164 of those were vaccinated). As such, it offers the potential to assess effects of the vaccine that may not be apparent in the short-term. Yet, this non-standard approach increases the potential for bias and given the disparity between the study’s findings and those of Simpson et al. [4], careful appraisal of the study’s design and analysis is warranted to establish whether the findings are robust.

Examination of the study by Phillips et al. [2], is also pivotal as the study’s findings were the foundation upon which vaccination of koalas with Klavax^TM^ is being pursued as a intervention to reduce rates of chlamydiosis in koala populations [3]. Risk vs benefit assessment is central to all medical and veterinary interventions. This is of heightened importance for prophylactic vaccines, since they are administered to otherwise healthy people or animals. Further, understanding the risks and benefits of a vaccine are of even greater importance when it is administered to an endangered species. In the case of the koala chlamydia vaccine, there has been no suggestion in the literature that it possesses a risk to koala safety. However, it remains imperative that any benefit of the vaccine be robustly established to justify its continued advancement and implementation.

Given the importance of the findings reported by Phillips et al., [2], here we report an independent analysis performed on the processed data collated and analysed by Phillips et al., [2], and highlight sources of inherent bias which have impacted the validity of the vaccine’s reported protective efficacy.

## Methods and Results

### Data description

Supplementary Table 3 from Phillips et al. [2], was downloaded and used as the basis of this analysis. The dataset includes records of 680 individual animals. 164 of these animals were recorded as being vaccinated, of which 49 were vaccinated at the time of the first record. As records of individual animals before vaccination were also included with non-vaccinated animals in the analysis of Phillips et al. [2], they reported records of 631 control animals (516 non-vaccinated and 115 pre-vaccination). However, as the disease and health status of koalas at the time of vaccination also provides information on pre-vaccination health, we also include the 49 koalas vaccinated at first record in the control group. As such, the total sample size for the control group in our re-analysis is 680 koalas (Table 1). Phillips et al. [2], Supplementary Table 3 includes a total of 4,362 records representing each individual assessment performed on a koala within the combined population.

**Table 1.** Control and vaccinated koalas by published study and vaccine formulations.

| Publication | Control animals | Vaccinated animals | Vaccine |
| --- | --- | --- | --- |
| **Phillips et al., 2025 [2]** | **680\*** | **164\*\*** | **Multiple** |
| *Waugh et al., 2016 [5]* | 30 | 30 | 3 doses of 3MOMP+ISC |
| *Desclozeaux et al., 2017 [6]* | 21 | 21 | 1 dose of 3MOMP+TriAdj |
| *Khan et al., 2016 [7]* | 0 | 15 | 1 dose of 3MOMP+TriAdj (n=10)<br>3 doses of 3MOMP+ISC (n=5) |
| *Khan et al., 2016 [8]* | 10 | 10 | 3 doses of rMOMP (unknown adjuvant) |
| *Quigley et al., 2023 [9]* | 23 | 46 | 1 dose of MOMP peptides+TriAdj |
| *Monitored koalas not previously included in a vaccine trial and unaccounted for vaccinated koalas* | 596 | 42 | *MOMP (unknown adjuvant)* |

*\*Note. Phillips et al. [2] (version published in 2025) refers to 631 control animals, however that figure did not include 49 animals that received a vaccine during their first encounter, yet health status at that initial encounter should logically have been included within the control group.*
*\*\*Phillips et al. [2] (version published in 2025) refers to 163 vaccinated animals, however records for 164 vaccinated animals are listed in Phillips et al, [2] Supplementary Table 3*

### Vaccine Administration

Animals included within this dataset represent the aggregate of 5 separate published vaccine trials and a sixth study performed in 2022 in which 18 previously vaccinated koalas were boosted, which is yet to be independently published. Phillips et al. [2], therefore combines multiple different vaccine types, adjuvants and dosing regimen (Table 1). The dataset also includes long-term monitoring data on the population.

### Identification of multiple errors in how the dataset was coded for the survival and disease probability analysis

The survival probability analysis conducted by Phillips et al. [2] using the coxph function from the survival package in R [11, 12], fits a model to explain the survival (or disease status) of an animal using empirical records of animals during discrete intervals of time (e.g. 0 – 3 yrs of age). The response variable in the analysis is whether an animal died (or became diseased) during an interval and the explanatory variables are traits that apply to that animal during that interval (e.g. they were vaccinated during that time).

During interrogation of Phillips et al. [2] Supplementary Table 3, a major error was identified in how the data was coded for analysis. For each of the 164 vaccinated koalas Tstart (which indicates the age of the koala at the beginning of the interval) for their first record was entered as zero and Tstop (the age of the koala at the end of the interval) the age of vaccination. This meant that their survival and disease status from birth until the age of vaccination was included in the vaccinated group despite animals being unvaccinated during that period. This is a substantial error that affects the outcome of the study by incorrectly attributing vaccination to survival and disease status of koalas prior to vaccination. Further, as the koalas necessarily survived to the time of vaccination and the majority of vaccinated koalas were healthy at the time of vaccination (see below), inclusion of the pre-vaccination interval artificially inflates apparent survival and decreases disease in the vaccinated cohort.

Additionally, for koalas that were included in the control group pre-vaccination and then vaccinated, the koalas were included in the analysis twice between birth and vaccination; once in the control group and once in the vaccinated group. Reanalysis of the survival and disease probabilities according to the method of Phillips et al. [2] with the Tstart of the first vaccination record corrected to the Tstop (the age of the koala at the end of the interval) of the last pre vaccination record produced identical results to those reported in the 2024 version of Phillips et al. [2]. As such, it can be concluded that this error in coding was introduced in the correction to the paper published in 2025.

We also identified an additional minor error in how the data was coded. Koala identity was included in the analysis of Phillips et al. [2] as a grouping or clustering variable to account for repeated measures of the same animal. However, koalas were entered as different individuals in the analysis when they were vaccinated, which represents a pseudoreplication error that artificially inflates statistical power as the records from the same koalas pre and post vaccination are not independent.

We corrected for these errors by altering the data coding as described above (Supplementary File 1; see supplementary information for detailed explanation of changes made).

### Reanalysis of survival and disease probabilities using the error corrected full dataset

Given the identified errors in the coding of the data in Phillips et al. [2] Supplementary Table 3 as described above, we refitted the survival and disease probability models using the coxph function from the survival package in R [11, 12] as per Phillips et al. [2], with those errors corrected (Supplementary File 1). These models include vaccination status, sex and year as explanatory variables. It should also be noted that Phillips et al [2] refers to 110 koala deaths within the population, 59 of which died with disease. However, Phillips et al [2] Supplementary Table 3 only contains records for 42 of 59 animals that died with disease and 0 of 61 deaths which occurred in the absence of disease. We reached out to the authors of Phillips et al. [2], to request these missing details but no further data was provided. We were therefore only able to include the 42 recorded deaths within our reanalysis, which correspond to those used by Phillips et al. [2] in their disease and survival probability analysis. Reanalysis of the corrected dataset showed no significant effect of vaccination status on either koala survival (p = 0.202; Figure 1) or disease (p = 0.798; Figure 2).

**Figure 1.**
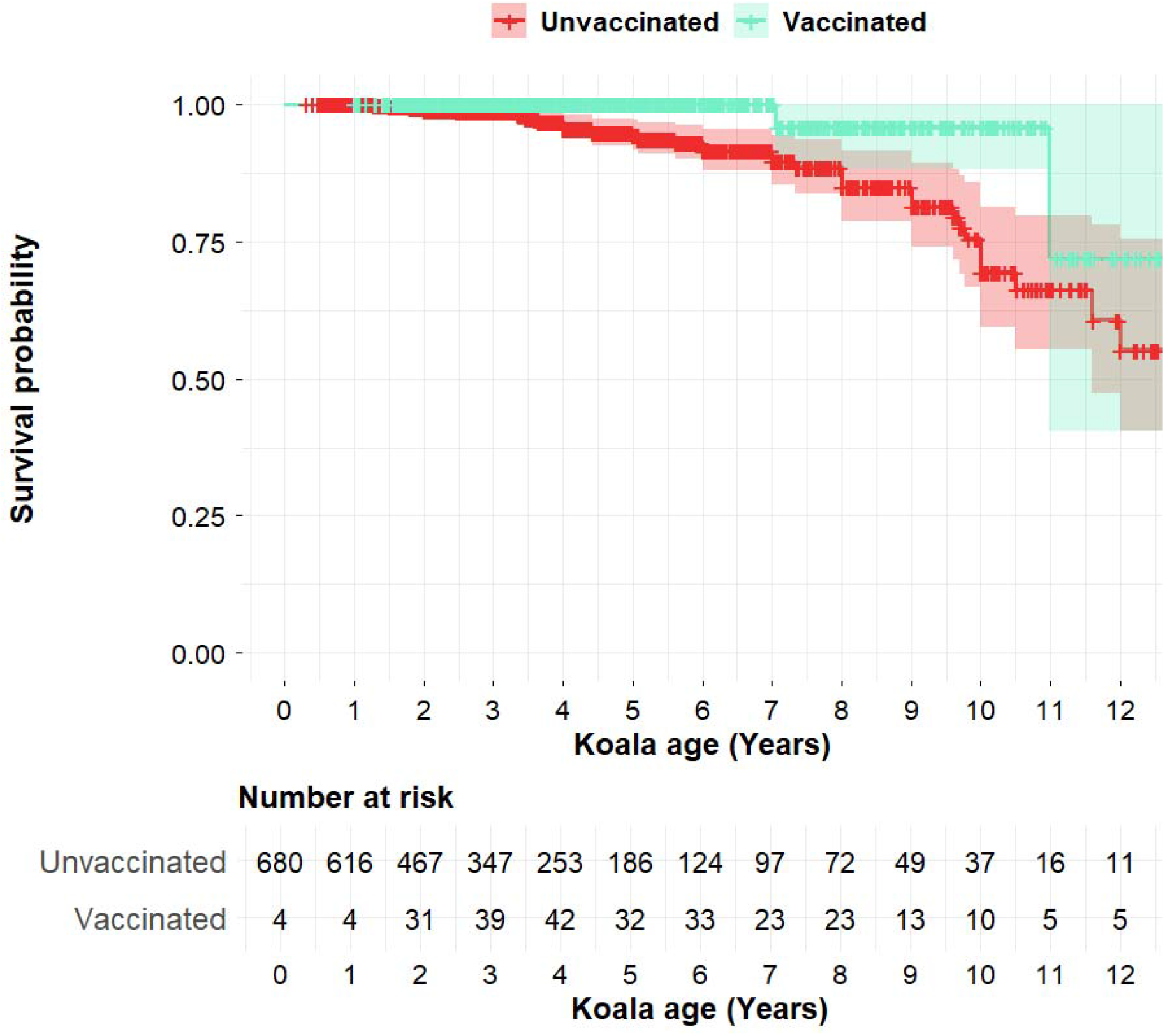
Fitted survival probability plots for unvaccinated and vaccinated koalas produced using the servminer package in R [13] based on the corrected data from Phillips et al. [2], Supplementary Table 3. The number at risk table indicates the number of records for koalas in each cohort at a given age.

**Figure 2.**
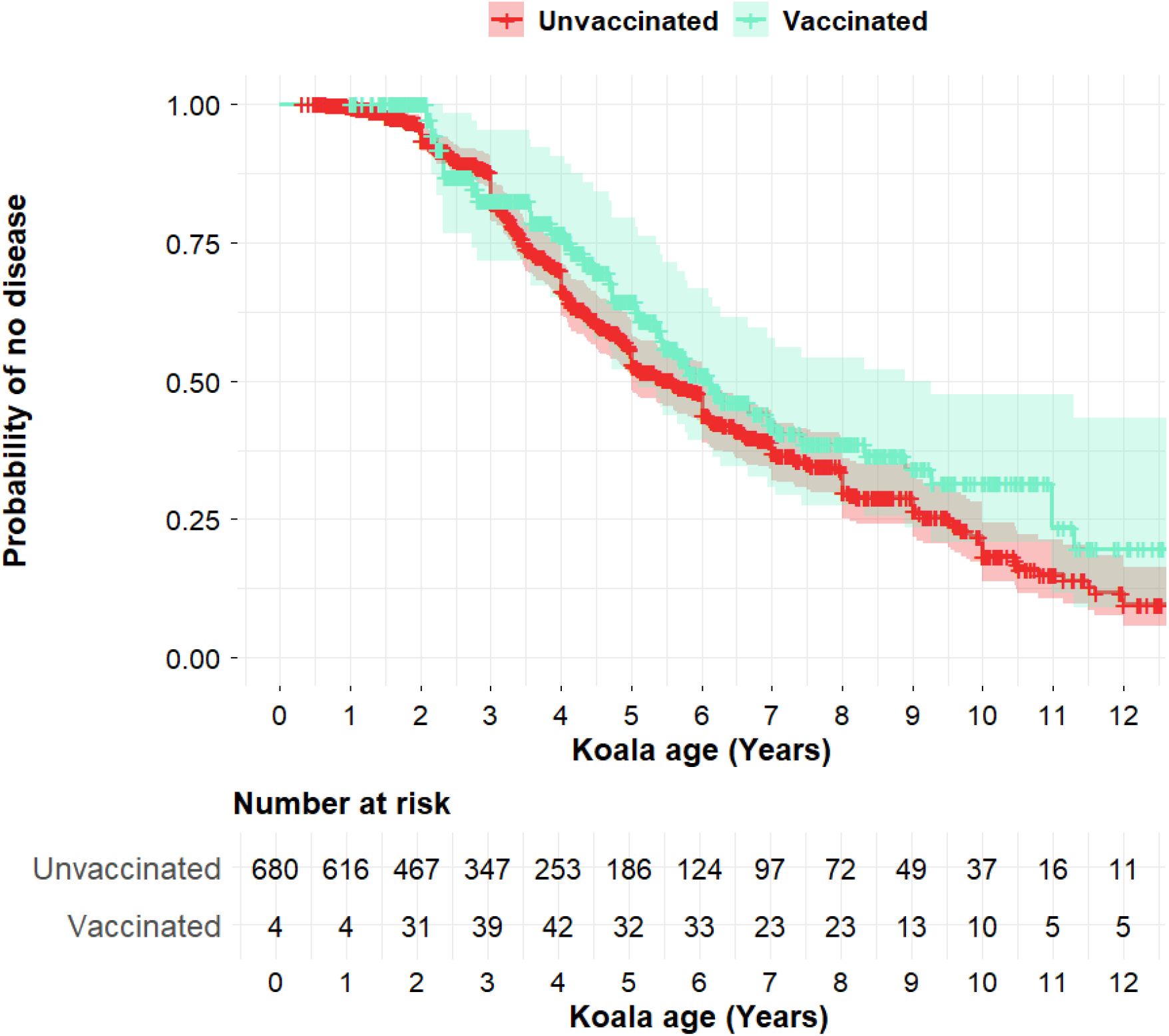
Fitted plots for the probability of no disease for unvaccinated and vaccinated koalas produced using the servminer package in R [13] based on the corrected data from Phillips et al. [2], Supplementary Table 3. The number at risk table indicates the number of records for koalas in each cohort at a given age.

### Inclusion of koalas with disease at first encounter introduces bias between the control and treatment groups

Of the 680 koalas in the dataset, 100 were recorded as having disease at first encounter and 21 were recorded as dying with disease at first encounter. It can be logically assumed for those 21 animals that a veterinary assessment performed at the first encounter deemed that they were too diseased to be released back into the wild and were euthanized on humanitarian grounds. The inclusion of these koalas in the control group in the original analysis of Phillips et al. [2] represents a significant bias in the dataset, as no koala presenting in such poor condition was vaccinated. Further, 100/680 (14.7%) of control koalas were diseased on first encounter, while only 4/164 (2.4%) koalas were diseased when they were vaccinated (Table 2). This represents a further bias in the analysis as the rate of disease at the beginning of records for each group was 6.1 times higher for the control group compared to the vaccinated group.

**Table 2.** Disease and death rates for Control and Vaccinated koalas.

|  | Koalas | Any Disease | Deaths | Disease at Tstart | Deaths at Tstart |
| --- | --- | --- | --- | --- | --- |
| Unvaccinated | 680 | 163 | 39 | 100 | 21 |
| Vaccinated | 164 | 26 | 3 | 4 | 0 |

### Corrected analysis with matched healthy control population

To remove the bias in the dataset introduced by including koalas that were diseased at first encounter/vaccination, we removed these individuals from the corrected dataset. Further, as koalas could only be included in the control group between birth and first encounter (no koala was vaccinated at birth), this interval was removed from the analysis for all koalas to ensure that the treatment and control groups were equivalent. This filtering also meant that koalas that were only encountered once were removed from the dataset. The resulting filtered dataset (Supplementary File 2) includes 439 control animals and 143 vaccinated animals, which represented 87% of vaccinated animals and 65% of control animals. This filtered dataset includes 3,302 health records were available for these animals, representing 76% of the total 4,362 records available in Phillips et al. [2].

A power calculation was performed based on a disease incidence rate in the control population of 15%. A sample size of 439 control and 143 vaccinated animals was predicted to have an 80% chance of detecting a vaccine efficacy of 64% at p=0.05.

Despite sufficient sample size to detect moderate or greater vaccine efficacy, the rate of disease in both vaccinated and control groups in the filtered dataset were almost identical (14.4% vs 12.6%). Furthermore, equivalent rates of death with associated disease were found in the control and vaccine group (2.5% vs 2.1%). Vaccine efficacy against disease was estimated to be only 12.3% (95%CI = -43.0% to 46.2%), while vaccine efficacy against death with associated disease was 16.3% (95% CI = -195.9% to 76.3%).

We also included within our analysis whether there was any difference in disease that was resolved over the course of the follow up period (i.e. cleared by last encounter), or unresolved disease that remained detectable at the time of the last record (Table 3). Surprisingly, disease was significantly more likely to remain unresolved in vaccinated animals; 13 of 143 (9.1%) vaccinated animals had unresolved disease compared with 27 of 439 (6.2%) of control animals (χ^2^: p =0.028 based on 81 cases of disease).

**Table 3.** Disease and death rates for Control and Vaccinated koalas after koalas with disease at first encounter/vaccination were removed from the dataset.

|  | Animals | Any Disease | Resolved disease | Unresolved disease | Deaths |
| --- | --- | --- | --- | --- | --- |
| Control | 439 | 63 (14.4%) | 36 (8.2%) | 27 (6.2%) | 11 (2.5%) |
| Vaccinated | 143 | 18 (12.6%) | 5 (3.5%) | 13 (9.1%) | 3 (2.1%) |

Further when we refitted the survival and disease probability models using the coxph function from the survival package in R [11, 12] as per Phillips et al. [2] using the filtered dataset, there was no statistically significant difference in either survival or rates of disease between the vaccinated and control groups (survival: p = 0.394; disease: p = 0.426; Figures 3-4).

**Figure 3.**
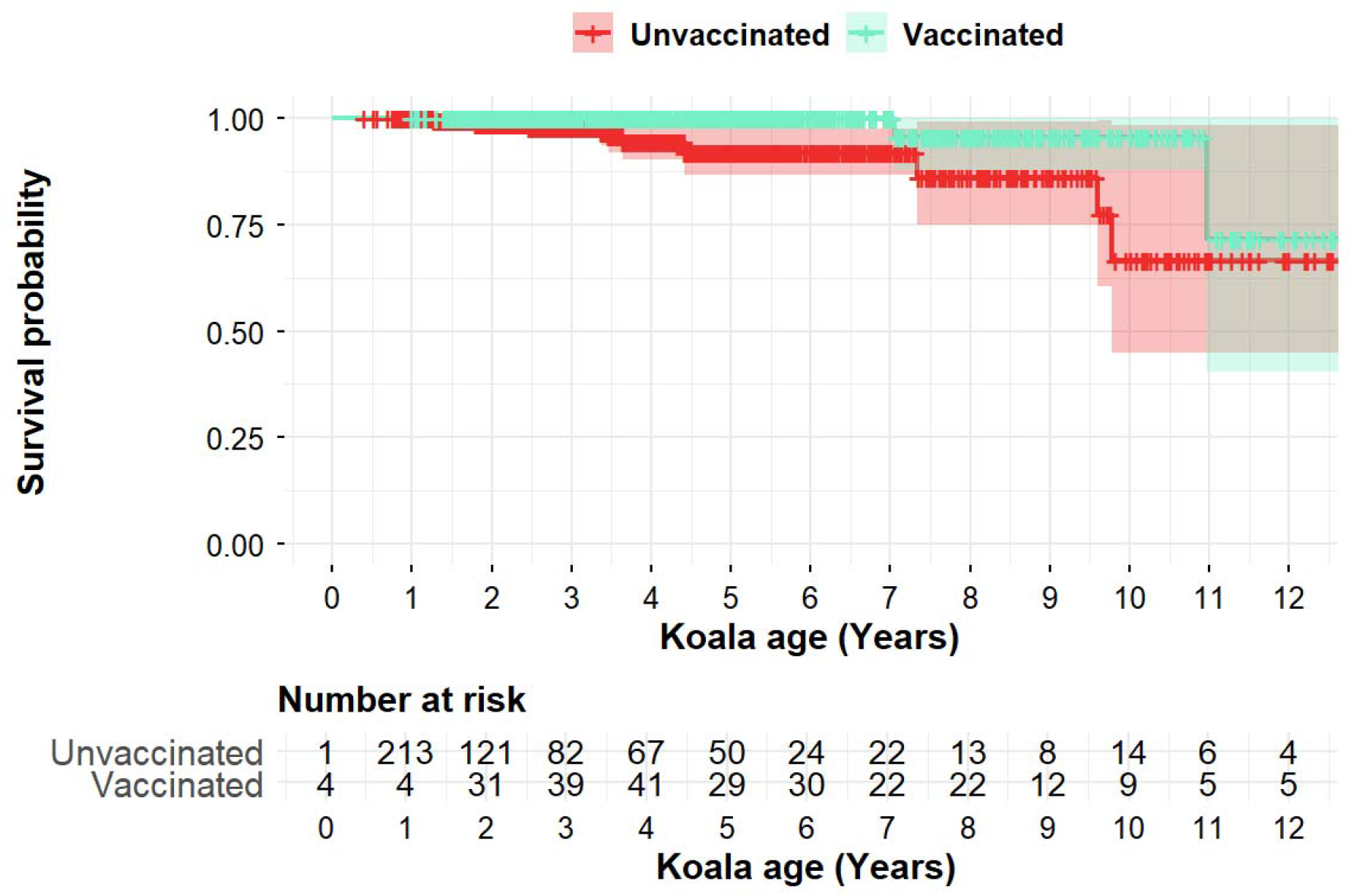
Fitted survival probability plots for unvaccinated and vaccinated koalas produced using the servminer package in R [13] based on the filtered data from Phillips et al. [2], Supplementary Table 3 where koalas with disease at first encounter/vaccination and the birth to first encounter interval removed. The number at risk table indicates the number of records for koalas in each cohort at a given age

**Figure 4.**
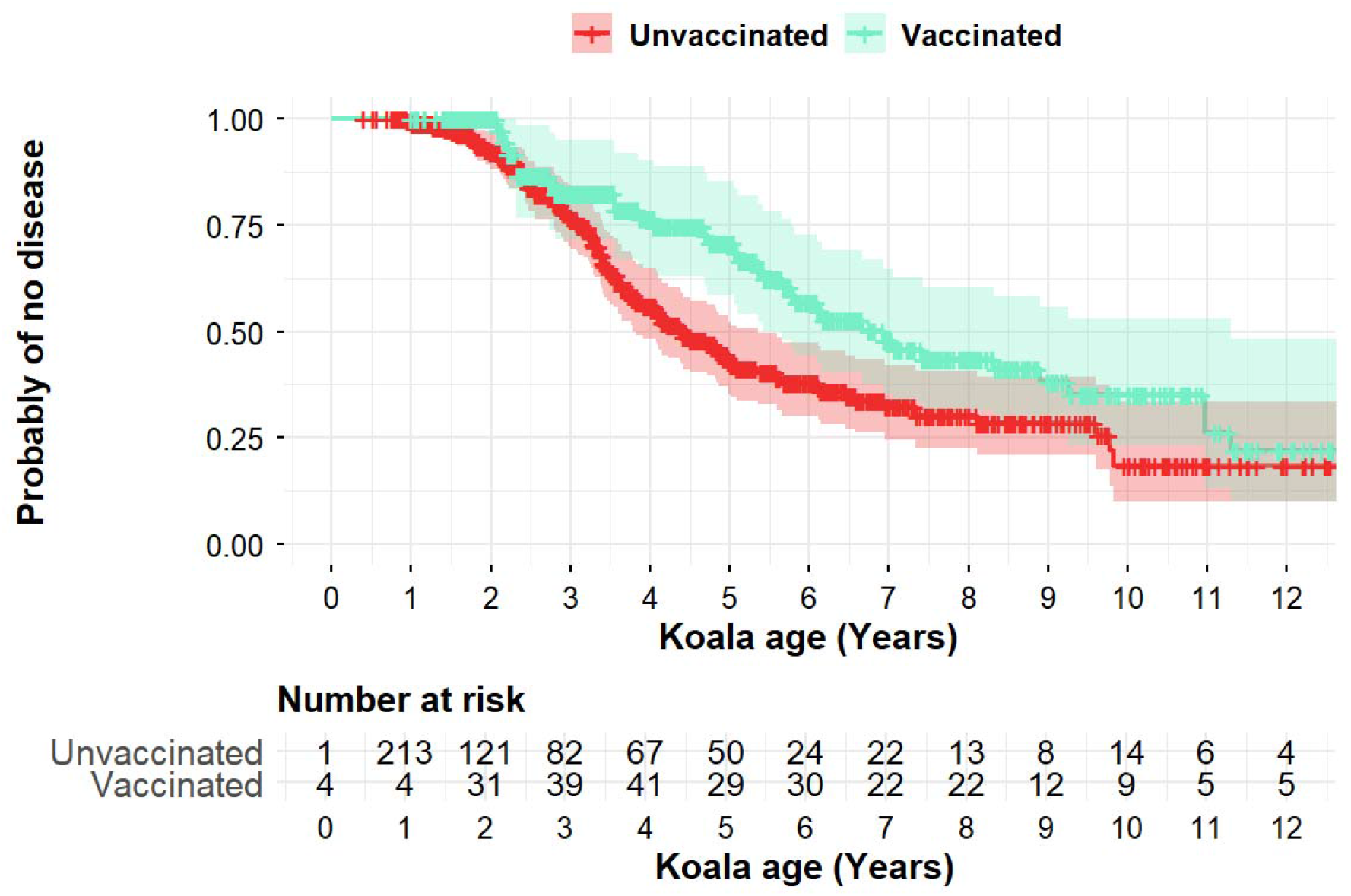
Fitted plots for the probability of no disease for unvaccinated and vaccinated koalas produced using the servminer package in R [13] based on the filtered data from Phillips et al. [2], Supplementary Table 3 where koalas with disease at first encounter/vaccination and the birth to first encounter interval removed. The number at risk table indicates the number of records for koalas in each cohort at a given age.

### Additional sources of bias in the dataset

Given the finding that vaccination has no statistically significant effect on disease risk and was associated with increased risk of unresolved disease, we next looked for any additional differences between the cohorts that could further bias the data. Within the filtered dataset (where diseased koalas on first encounter were removed; Supplementary File 2) we first compared koala age and how long the koalas were tracked for (Table 4). Vaccinated koalas were on average slightly older than the control group at the time of first vaccination/first encounter and also at the time of the last record (Table 4). This is to be expected as most of the vaccinated animals began in the control group prior to vaccination. Consistent with these averages, a higher proportion of records were collected from control animals at a younger age compared to the koalas when vaccinated (Figure 5A).

**Table 4.** Age of koalas at first and last encounter.

|  | Animals | Average Age at first encounter | Average Age at last encounter | Average duration |
| --- | --- | --- | --- | --- |
| Control | 439 | 2.1 yoa | 3.5 yoa | 1.4 years |
| Vaccinated | 143 | 3.5 yoa | 5.3 yoa | 1.8 years |

**Figure 5.**
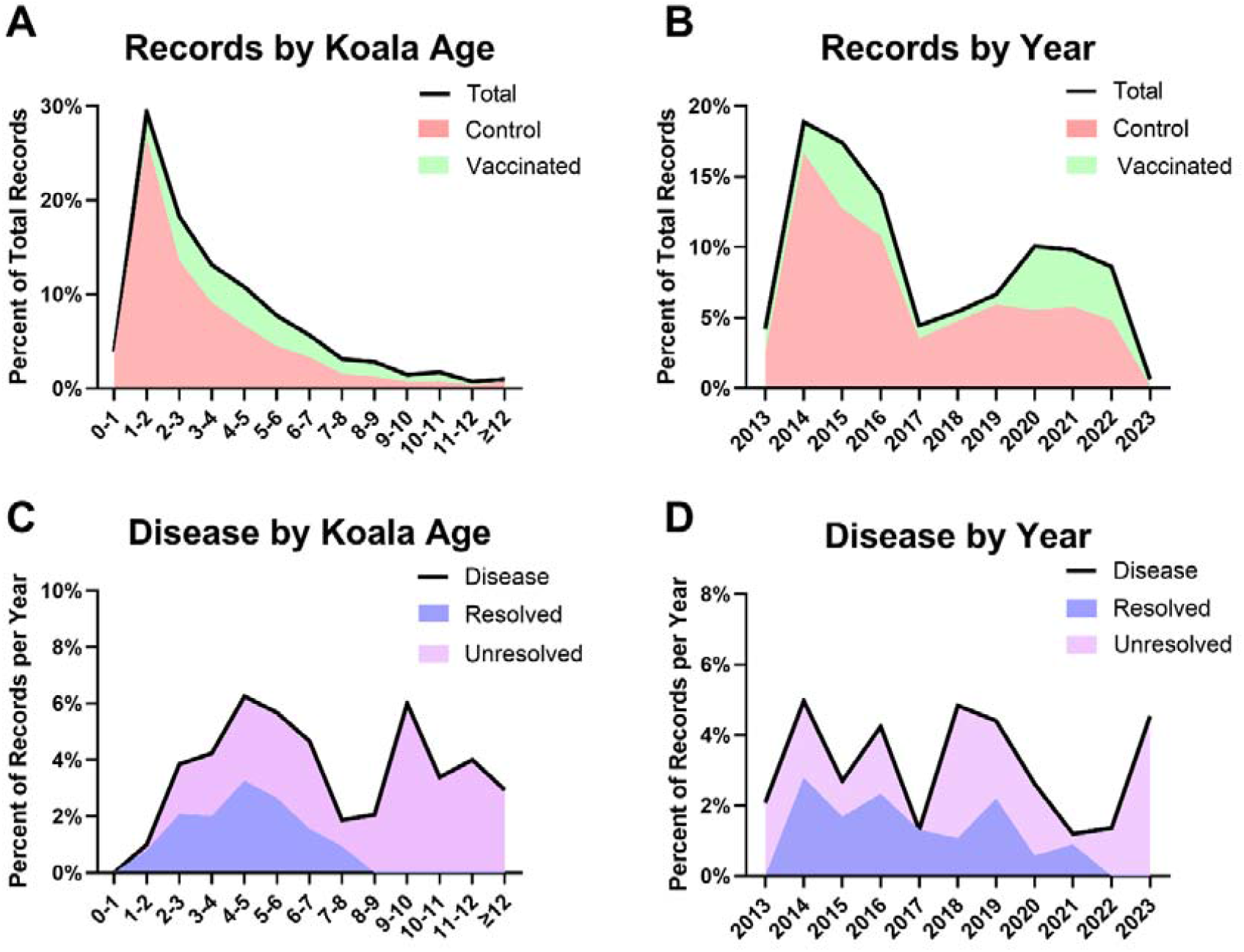
A/B. Number of records included within the filtered dataset separated based on koala age or year of in which koala health assessment was performed. **C/D**. Percent of records in which disease was reported separated based on koala age or year in which koala health assessment was performed.

Separation of records based on year of study showed two peaks, during which most observations occurred, 2014-2016 and 2020-2022. A larger proportion of records from control animals came from 2014-2016 and a larger proportion of records from vaccinated animals came during the period 2020-2022 (Figure 5B).

Overall disease was fairly evenly distributed throughout the life of the koala (Figure 5C). Although rates of disease were lowest between the ages of 0-2, this is likely due to younger koalas having less exposure to *C. pecorum* having not reached sexual maturity but also a result of filtering the dataset to only include animals that were healthy at first encounter. Disease was more likely to be resolved for koalas at a younger age. Over half of the incidences of disease were resolved for koalas between the age of 1 and 6, whereas no incidences of resolved disease were identified in koalas over the age of 8 (Figure 5C).

The distribution of disease was relatively uniform over time, with typically between 2 and 5% of records listing the presence of disease (Figure 5D). There was also no obvious trend comparing resolved vs unresolved disease, except for the final year of the study, which was all unresolved as no further records were available.

When the distribution of disease by koala age and year of study was considered for the vaccinated and control koala groups separately, it was evident that unresolved disease was more frequent in vaccinated koalas at a younger age (Figure S3). In vaccinated koalas less than 6 years of age, 15 of 124 (12.1%) had recorded disease, of which 5 (33%) were resolved, compared to 58 of 400 (14.5%) control animals under 6 years of age that had disease recorded, of which 35 of 58 (60%) were resolved (Table S1). The age difference between vaccinated and control animals is therefore unable to explain the higher proportion of unresolved disease in vaccinated animals. There was also a spike in unresolved disease in vaccinated koalas between 2018 and 2019, however this is likely an artifact of low sample numbers collected during this period (Figure S3).

## Discussion

Chlamydia infection is a frequent cause of disease amongst koala populations throughout the Australian States of Queensland and New South Wales [14] and has been linked to population decline and collapse in some areas [15]. Additionally, approximately half of koalas admitted to wildlife hospitals present with chlamydial disease, which causes cystitis, reproductive pathology and/or conjunctivitis, often progressing to infertility and/or mortality [16, 17]. Thus, safe and effective interventions for chlamydia are greatly needed to respond to declining koala populations and improve koala welfare. However, the risks and benefits of any such intervention need to be carefully assessed. In this study, we examined and reanalysed the data and findings of Phillips et al. [2], which provided the primary evidence for the chlamydia vaccine, Klavax^TM^ (Tréidlia Ltd), to be approved for minor use by APVMA. Our analysis revealed a fundamental flaw in the way the data was encoded and analysed as well as a stratification bias that led to the erroneous finding that the vaccine provided protection from disease and disease associated death.

Upon identification of the coding error, we contacted both the authors and publishing journal of Phillips et al., [2] and we have been informed that a correction is in preparation. However, here we have also provided an analysis of the corrected dataset using the same method as Phillips et al. [2] as it was necessary before any further reanalysis of the data could take place. This corrected analysis showed a small but non-statistically significant trend for vaccination to provide protection from disease and death with disease.

After correction for the coding error, our reanalysis of the data revealed a clear stratification bias whereby koalas were approximately six-times more likely to have disease at the first timepoint (100 of 680 control compared to just 4 of 164 vaccinated koalas) [18]. To exclude this source of bias from the study, a filtered dataset was produced that included 439 control and 143 vaccinated animals, all of which were healthy at the first assessment/vaccination and had their health monitored during a follow up period. Despite the reduced sample size of the filtered dataset, it remained the largest koala chlamydia vaccination dataset analysed to date and a power analysis showed it retained the power to detect a moderate or greater vaccine efficacy. Unfortunately, following the removal of the stratification bias, there was no difference in the frequency of disease or disease associated mortality between the control and vaccination cohorts; 12.6% of vaccinated animals contracted disease and 2.1% died compared to 14.4% of control animals that contracted disease and 2.5% that died.

Surprisingly, vaccinated koalas appeared to be less likely to return to health once they were recorded as having disease compared with control koalas. Only 5 of 18 vaccinated koalas were cleared of disease by their last health assessment compared to 35 of 63 koalas within the control group. If taken at face value such an observation may be alarming. Vaccinated koalas were slightly older than control koalas and therefore may have had less opportunity to clear disease; however, this effect is likely minimal as the average age of first encounter for both groups was younger than the average age of disease onset. Nevertheless, there may be other inherent differences between the cohorts such as if or when veterinary care/intervention was provided that may explain the higher rate of unresolved disease in the vaccinated cohort. As such, we would caution against any assumption that this observation represents a safety signal.

Overall, the findings of this independent analysis should serve to emphasize the importance of performing randomised, observer blinded, placebo-controlled methods for the assessment of veterinary interventions including vaccinations. Such methodologies are standard practice during the development of human vaccines and are recommended by the APVMA. In agreement with our reanalysis of the data from Phillips et al., [2] after correcting for inherent bias in the study design, when the same vaccine was assessed within a randomised, observer blinded, placebo-controlled trial there was no evidence of a protective effect of vaccination [4]. This was despite a moderate sample size of 102 vaccinated wild koalas and 94 placebo control koalas that would have been sufficient to detect moderate to high levels of efficacy [4]. However, this undesirable result appears to have been largely ignored, while the favourable results of the flawed study, reanalysed here, have been promoted [2, 3].

Considering the reanalysis presented here and the findings of Simpson et al. [4], both indicating no protective benefit for koalas, the general use of this vaccine formulation by veterinarians to prevent or reduce chlamydial disease in koalas should not be recommended. Instead, we advocate for the continued development and testing of an improved vaccine formulation and the advancement of alternative intervention strategies. Our reanalysis also highlights that attempting to assess vaccine efficacy using uncontrolled or inappropriately controlled field data is inherently open to errors in processing and systemic biases due to its complex structure. By contrast, the study by Simpson et al., [4] demonstrates that randomised, observer blinded, placebo-controlled studies are possible in wild, free ranging koalas. Therefore, we argue that further efforts to develop and test chlamydia vaccines for koalas should utilize this controlled design to avoid the introduction of bias and ensure robust valid findings.

## References

1. Department of Agriculture, W.a.t.E., Australian Federal Government, Conservation Advice for Phascolarctos cinereus (Koala) combined populations of Queensland, New South Wales and the Australian Capital Territory, W.a.t.E. Department of Agriculture, Australian Federal Government, Editor. 2022, Department of Agriculture, Water and the Environment: Canberra.

2. Phillips, S., et al., Author Correction: Immunisation of koalas against Chlamydia pecorum results in significant protection against chlamydial disease and mortality. NPJ Vaccines, 2026. 11(1).

3. Pollak, N.M., S. Phillips, and P. Timms, Koalas first: lessons from a wildlife Chlamydia vaccine. Trends Microbiol, 2026. 34(6): p. 601–613.

4. Simpson, S.J., et al., Evaluation of Chlamydia pecorum major outer membrane protein vaccine a management tool in wild koala (Phascolarctos cinereus) populations. Sci Rep, 2025. 15(1): p. 30601.

5. Waugh, C., et al., A Prototype Recombinant-Protein Based Chlamydia pecorum Vaccine Results in Reduced Chlamydial Burden and Less Clinical Disease in Free-Ranging Koalas (Phascolarctos cinereus). PLoS One, 2016. 11(1): p. e0146934.

6. Desclozeaux, M., et al., Immunization of a wild koala population with a recombinant Chlamydia pecorum Major Outer Membrane Protein (MOMP) or Polymorphic Membrane Protein (PMP) based vaccine: New insights into immune response, protection and clearance. PLoS One, 2017. 12(6): p. e0178786.

7. Khan, S.A., et al., Antibody and Cytokine Responses of Koalas (Phascolarctos cinereus) Vaccinated with Recombinant Chlamydial Major Outer Membrane Protein (MOMP) with Two Different Adjuvants. PLoS One, 2016. 11(5): p. e0156094.

8. Khan, S.A., et al., Humoral immune responses in koalas (Phascolarctos cinereus) either naturally infected with Chlamydia pecorum or following administration of a recombinant chlamydial major outer membrane protein vaccine. Vaccine, 2016. 34(6): p. 775–82.

9. Quigley, B.L., et al., Reduction of Chlamydia pecorum and Koala Retrovirus subtype B expression in wild koalas vaccinated with novel peptide and peptide/recombinant protein formulations. Vaccine X, 2023. 14: p. 100329.

10. Australian_Pesticides_and_Veterinary_Medicines_Authority, Guideline for the registration of new veterinary vaccines. 2020: https://www.apvma.gov.au/registrations-and-permits/data-guidelines/veterinary-data-guidelines/specific-guidelines/new-vaccine.

11. Therneau, T., A package for survival analysis in R. R package version, 2015. 2(7): p. 2014.

12. R-Core-Team, R: A language and environment for statistical computing. 2012, R Foundation for Statistical Computing: Vienna, Austria.

13. Kassambara, A., M. Kosinski, and P. Biecek, survminer: Drawing Survival Curves using’ggplot2’. CRAN: Contributed Packages, 2016.

14. Robbins, A., et al., Longitudinal study of wild koalas (Phascolarctos cinereus) reveals chlamydial disease progression in two thirds of infected animals. Scientific reports, 2019. 9(1): p. 1–9.

15. McAlpine, C., et al., Conserving koalas: a review of the contrasting regional trends, outlooks and policy challenges. Biological Conservation, 2015. 192: p. 226–236.

16. Gonzalez-Astudillo, V., et al., Decline causes of Koalas in South East Queensland, Australia: a 17-year retrospective study of mortality and morbidity. Scientific reports, 2017. 7: p. 42587.

17. Kerlin, D.H., L.F. Grogan, and H.I. McCallum, Insights and inferences on koala conservation from records of koalas arriving to care in South East Queensland. Wildlife Research, 2022. 50(1): p. 57–67.

18. Banack, H.R., et al., Collider Stratification Bias I: Principles and Structure. Am J Epidemiol, 2024. 193(2): p. 238–240.

