## Supplementary Information for "Reanalysis of data from ten years of koala chlamydia vaccine trials reveals inherent bias and a lack of protective efficacy"

**Supplementary information on data coding error and correction.**

Interrogation of Phillips et al. [2] Supplementary Table 3 identified 49 individual koalas that were vaccinated on first encounter (Example: Arya - Figure S1). For each of these animals Tstart (which indicates the age of the koala at the beginning of the interval) was entered as zero and Tstop the age of vaccination. This meant that their survival and disease status from birth until the age of vaccination was included in the vaccinated group even though the animal was unvaccinated during this period. To correct these 49 records, a new column was added to the spread sheet ‘Vx_corrected’ where the vaccination status (Vx) was corrected from ‘1’ to ‘0’ to indicate the animal was unvaccinated during that period (Figure S1).

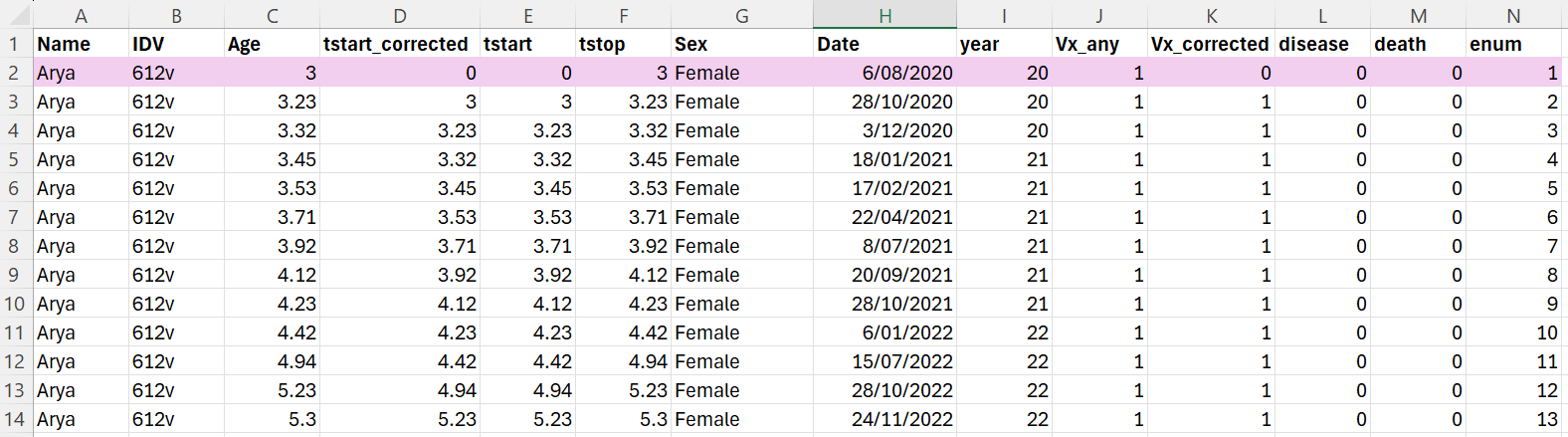

**Figure S1.** Example record for koala included in control group and then vaccinated

A further 115 individual koalas were first included in the control group pre-vaccination and then vaccinated (Example: Adder - Figure S2). For each of these animals the Tstart of their first vaccinated record was entered as zero and Tstop the age of vaccination. As with the example above, this meant that their survival and disease status from birth until the age of vaccination was included in the vaccinated group, however it was also included a second time in the control group. To correct this error, a new column was added to the spread sheet ‘Tstart_corrected’, where the Tstart values were changed to the Tstop values of the previous record, so that the period was only covered once. Additionally as with the koalas vaccinated at first encounter, the vaccination status (Vx) was corrected from ‘1’ to ‘0’ to indicate the animals were unvaccinated during that period (Figure S2).

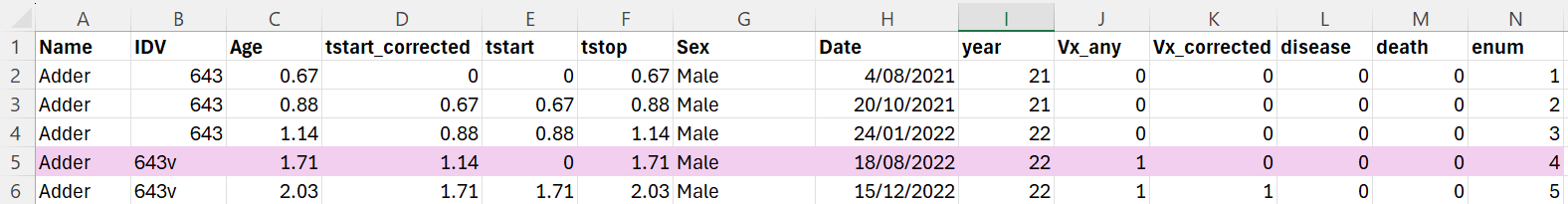

**Figure S2.** Example record for koala included in control group and then vaccinated

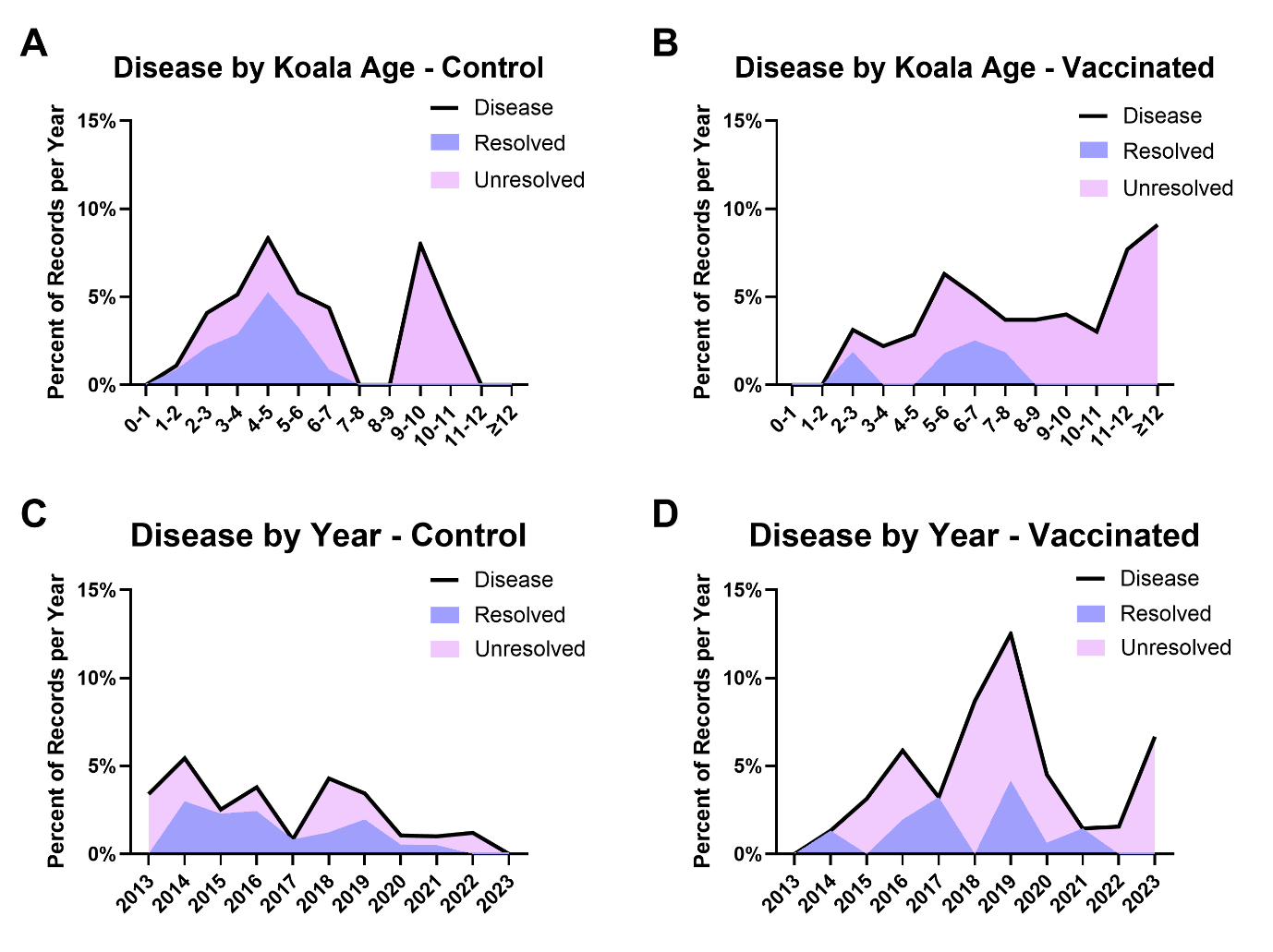

**Figure S3. A-D.** Percent of records from control and vaccinated groups in which disease was reported separated based on koala age or year of in which koala health assessment was performed. Full data set provided in supplementary file 2.

**Table S1:** Disease and death rates for Control and Vaccinated koalas under 6 years of age

|  | Animals | Any Disease | Resolved disease | Unresolved disease | Deaths |
| --- | --- | --- | --- | --- | --- |
| Control | 400 | 58 (14.5%) | 35 (8.8%) | 23 (5.8%) | 8 (2.0%) |
| Vaccinated | 124 | 15 (12.1%) | 5 (4.0%) | 10 (8.1%) | 3 (2.4%) |
